# The Case For and Against Double-blind Reviews

**DOI:** 10.1101/495465

**Authors:** Amelia R. Cox, Robert Mongomerie

**Affiliations:** Biology, Queen’s University, Kingston, Ontario, Canada

## Abstract

To date, the majority of authors on scientific publications have been men. While much of this gender bias can be explained by historic sexism and discrimination, there is concern that women may still be disadvantaged by the peer review process if reviewers’ unconscious biases lead them to reject publications with female authors more often. One potential solution to this perceived gender bias in the reviewing process is for journals to adopt double-blind reviews whereby neither the authors nor the reviewers are aware of each other’s identities and genders. To test the efficacy of double-blind reviews, we assigned gender to every authorship of every paper published in 5 different journals with different peer review processes (double-blind vs. single blind) and subject matter (birds vs. behavioral ecology) from 2010-2018 (n = 4865 papers). While female authorships comprised only 35% of the total, the double-blind journal *Behavioral Ecology* did not have more female authorships than its single-blind counterparts. Interestingly, the incidence of female authorship is higher at behavioral ecology journals (*Behavioral Ecology* and *Behavioral Ecology and Sociobiology*) than in the ornithology journals (*Auk, Condor, Ibis*), for papers on all topics as well as those on birds. These analyses suggest that double-blind review does not currently increase the incidence of female authorship in the journals studied here. We conclude, at least for these journals, that double-blind review does not benefit female authors and may, in the long run, be detrimental.

## Introduction

For the past 25 years, there has been a welcome flurry of interest in the role and relative success of women in the process of scientific publication (e.g., Gilbert, Williams & Lundberg, 1994; Tregenza, 2002; Budden et al., 2008; Cho et al., 2014). The main foci of this research have been to assess the contributions of women to authorship, editorship, and collaborations, as well as to determine whether manuscript reviewers might be biased with respect to the gender, nationality, and reputation of authors. In a global, multidisciplinary, bibliometric analysis of 5.5 million academic papers published from 2008 to 2012, for example, Larivière et al. (2013) found that women published relatively fewer papers than men, were less likely to be first or last author on multi-authored papers, and, even when women were in these ‘dominant author’ positions, their papers were less likely to be cited than when men were first or last author. These various gender gaps varied by discipline, and author nationality but are echoed in a recent analysis of both manuscript submissions and published papers in 7 ecology journals (Fox, Ritchey & Paine, 2018). Several studies indicate that this gap has been ameliorating over the most recent decade, suggesting that changes in society at large, and in the scientific publishing process, in particular, are proving to be beneficial to female academics.

While it is unclear whether—but expected that—gender biases against women will influence research careers (Larivière et al., 2013), factors that reduce publication rate and quality will certainly have a negative impact. For that reason, many journals have adopted a double-blind reviewing policy wherein the reviewers are not revealed to the authors, and anything that might identify an author is removed from the manuscript before review. While the reasons for adopting double-blind reviews are laudable, there are some costs (see Discussion) and, to date, there is largely controversial evidence that such policies are having the desired effect. For instance, Budden et al. (2008) found that female first authorship was 7.9% higher in *Behavioral Ecology* after that journal switched from single-blind to double-blind reviews, while five comparable ecology journals that retained single-blind reviews showed no increase in the incidence of female authorship. However, others have suggested that different statistical analyses would have been more appropriate and have shown that the incidence of female authorship has steadily increased across all journals, regardless of peer review style (Engqvist & Frommen, 2008; Webb, Hara & Freckleton, 2008).

In the present study, we tested the idea that double-blind reviews have influenced the publication success of female authors. We considered three possible approaches to such a study. First, real manuscripts submitted to a given journal could be sent to typical reviewers in a paired design where one reviewer sees the author details, and the other does not (e.g., Tomkins, Zhang & Heavlin, 2017). Alternatively, author names could be fictitious but readily identifiable as either male or female, again in a paired design. This may be the most powerful experimental method, but it requires a considerable contribution from a journal and would need to be run for several issues or even years to generate a large enough sample for analysis.

Second, real or fake manuscripts can be assigned randomly to multiple readers to assess the effects of different author-gender combinations on perceived quality (e.g., Borsuk et al., 2009; Knobloch-Westerwick, Glynn & Huge, 2013; Okike et al., 2016). This method is excellent with respect to experimental design as so many potentially confounding factors can be controlled but it requires a fairly large number of willing and knowledgeable readers. Typical reviewers are unlikely to be willing to devote time to such an experiment, so this sort of study usually employs student readers. As a result, the subject matter in the papers used in such experiments is often kept fairly general, and the results may not reflect the responses of expert reviewers to field-specific manuscripts.

Third, a study can assess the differences between papers published in journals with and without—or in the same journal before and after (e.g., Budden et al., 2008)—it adopts double-blind reviews. This method has the advantage of involving large numbers of readily accessible papers, and, at least for comparisons between journals, can reveal trends over a period of years. The disadvantages are that submission and acceptance rates cannot be assessed, and different journals, even in the same field, might attract a different proportion of male and female authors, or submissions from different geographic regions, or with a different taxonomic or subject focus. Despite these limitations, we adopted this approach in the present study and attempted to control for differences between journals by comparing journals that we felt were very likely to attract the same authors and manuscripts, and by comparing publications that had the same taxonomic focus (birds) within those journals.

## Methods

### Data collection

We began this study to assess the potential advantages of two ornithological journals adopting a double-blind reviewing policy, *The Auk* (hereafter AUK) and *The Condor* (CONDOR), both now published by the American Ornithological Society. To do that, we compared recent publications (2010-2018) in those two journals to papers published in *Behavioral Ecology* (BE), a journal with double-blind reviews (since 2001) but similar journal impact factors (2017 IF = 2.44, 2.72, and 3.35, respectively). Only ~30% of the papers in BE in our dataset were about birds, so we also compared papers in BE with those in Behavioral Ecology and Sociobiology (BES), a single-blind journal with a similar audience and citation rate (2017 IF = 2.47) to BE. Because BE and BES had substantially more international authors than AUK and CONDOR, we added *The Ibis* (IBIS) to our analysis to see if author nationality might be important. IBIS uses single-blind reviews and is published by the British Ornithologists’ Union (2017 IF = 2.23).

For the 5445 papers published between 2010 and 2018 in those 5 journals, we assigned a gender to each authorship, noting the first and last authorships of each paper. We defined ‘authorship’ as each author on each paper; many authors publish multiple papers per year and thus account for multiple authorships. We assigned gender based solely on the perceived genders of first names rather than searching the internet for more information. Thus, we assumed that a reviewer would determine gender based on first names and would not have any additional information. For unfamiliar names, we used www.gpeters.com/names/baby-names to identify gender, requiring one gender to be >2x as likely as the other, otherwise, we scored it as ambiguous. For some papers authorships could not be assigned a gender because (i) only first initials were listed, (ii) the order of given and surnames was unclear (e.g., Asian names), or (iii) names were not consistently gendered (e.g., Robin which is only 1.53 times more likely to be male). In all, 580 papers with at least one authorship of ambiguous gender were excluded from all analyses, resulting in 4865 papers for the analyses presented here. Each paper was also scored as being about birds or other topics.

### Statistical analysis

We tested for gender biases in published papers, comparing journals and testing whether patterns changed over the 9 years in our sample. For all papers, we assessed the odds of having any female authorships in a paper and the proportion of authorships that were female. For single-author papers, we assessed the odds that the authorship was female. For multi-author papers, we assessed the odds of having a female authorship in the first or last position.

For each response variable, we performed a binomial logistic regression testing for associations between female authorships and journal, year, and their interaction. When testing for whether there were any female authorships on papers, we included the total number of authorships to account for the increase in female authorships as total authorships increases.

To test whether research collaborations lead by women had higher proportions of women involved as coauthors than collaborations lead by men, we looked for associations the proportion of female-authorships before the last authorship (i.e. collaborators) and the assumed gender of the last authorship, controlling for the journal, year, and their interaction.

We conducted all analyses using data from papers on all topics, as well as focusing only on papers about birds (see Table 1 for sample sizes). The vast majority of papers had <7 authors (95-96%; Fig. S1), so we also conducted analyses excluding all papers with >6 authorships. Including papers with long author lists did not affect the results (see Statistical Supplement S1 and S2).

For all analyses, we used a generalized linear model with binomial error and logit link function. Results are calculated as odds ratios (OR) then converted to percentages for ease of presentation (see Statistical Supplements for details). For all summary statistics, 95%CL are presented in square brackets. We report likelihood ratio chi-squares (LR χ^2^) for the variable of interest, testing the significance of removing that term from the model. To compare journals, we used Tukey posthoc tests on model results.

All analyses were conducted in R version 3.5.1 (R Core Team 2018). R scripts, analysis output, and raw data are deposited at Open Science Framework.

## Results

### Any female authorships

As expected, the odds of a paper having at least one female authorship increased with the total number of authors on the paper (Table 2, Fig. S2). The odds of a paper having at least one female authorship increased from 2010-2018 in AUK, CONDOR, IBIS, and BES but not at the double-blind BE (Fig. S3A), although the differences in slope are not significant (year*journal interaction, Table 2). As of 2018, BES has a higher percentage of papers with at least one female authorship (82% [78, 86]) than any other journal (BE 75% [70, 79], AUK 73% [66, 79], CONDOR 73% [65, 80], IBIS 72% [65, 79]) (Fig. S3C). The patterns were similar for papers specifically about birds (Table 2, Fig. S3B-C).

### Percentage of female authorships per issue

Across all papers (n = 4865) and years (n = 9), there were fewer female (mean 35%) than male (mean 65%) authorships (Fig. 1A), and this was true in almost every issue of all journals (n = 254 of 264 issues in 5 journals). Although the percentage of female authorships increased overall from 2010 to 2018, that rate differed significantly among journals (year * journal; Table 2; Fig. 1A). The percentage of female authorship in BES, AUK and IBIS increased significantly from 2010 to 2018 (per year by 4.3% [2.0, 6.6], 5.3% [2.0, 8.7], and 4.6% [1.0, 8.4], respectively). However, this was not the case for BE (0.1% [–2.1, 2.4]) or CONDOR (1.4 % [–2.0, 4.9]). Across all years BE and BES had a higher percentage of female authorships than any of the ornithology journals, and currently (2018) these differences are significant (Tukey posthoc tests, p < 0.05; see Statistical Supplement 1), except for the difference between BE and AUK (Fig. 1B).

**Figure 1:**
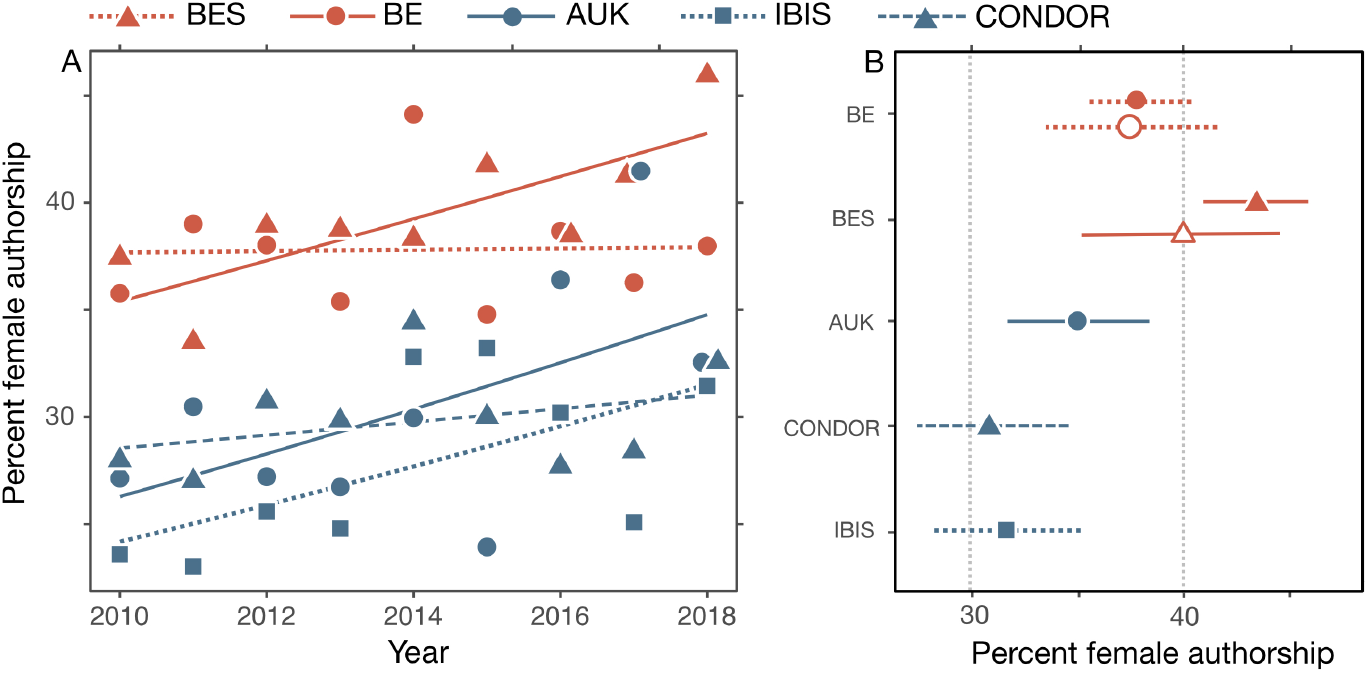
All female authorships in 5 journals from 2010 to 2018. (A) Female authorships as the percent of total authorships on all topics, with binomial trendlines. (B) Percentage of female authorship in 2018 (±95%CI) for each journal as well as for bird papers in BE and BES (open symbols). Percentages were calculated as marginal means of the models shown in Table 2. Papers with ambiguous authorships are not included. See Table 1 for sample sizes.

For papers about birds, the 2010-18 trends were similar to those for all papers (Fig. S4), but the rate of increase did not differ significantly across journals (year*journal interaction, Table 2). For papers published in 2018 only the differences between BES and both CONDOR and IBIS were significant (Tukey posthoc tests, p < 0.05; Fig. 1B).

### First-authorships

From 2010 to 2018, female first-authorships per year increased in all single-blind journals (BES 2.7% [−1.4, 7.2], AUK 7.8% [1.4, 14.6], CONDOR 4.4% [−2.3, 11.7], IBIS 6.3% [−0.5, 13.5]) but not in BE (−0.5% [−4.6, 3.8]), the only double blind journal in our study (Fig. 2A). These differences in the rate of change are not significant (year * journal, Table 2). In 2018, BES had the highest percentage of papers with female first-authorship (Fig. 2B), although all journals actually had higher (or comparable in the case of CONDOR) rates of female first authorship than the overall 2010-2018 percentage of female authorship in these journals (35%). BES had the highest percentage of female first-authorships in 7 of 9 years. Results were similar for bird-specific papers (Table 2, Fig. 2B, Fig. S5A).

### Last-authorships

The percentage of female last-authorships was generally stable or increasing slightly between 2010 and 2018 (per year, BE 2.4% [−2.1, 7.2], BES 3.9% [−0.9, 8.7], AUK 3.4% [−3.6, 11.0], CONDOR −1.5% [−8.7, 6.1], IBIS 5.0% [−3.2, 14.0]; Fig. 2C). Differences in the rate of increase across journals were not significant (journal*year, Table 2). In all journals, the percentage of female last-authorships was lower than for than female first-authorships, with IBIS having the lowest proportion of female last-authorships in 5 of the 9 years.

These differences between the behavioral ecology and ornithology journals seem to be driven by papers about non-bird taxa. Considering only papers about birds, last-authorships did not vary significantly among journals or years in our sample (Table 2, Fig. S5B).

In contrast to first-authorships, by 2018 all journals had lower percentages of female last-authorships than the overall percentage of female authorship (35%) in these journals (Fig. 2D). By 2018, all journals had comparable percentages of female last-authorships on bird papers (22%-27%) but the percentages of female last-authorships were higher in the behavioral ecology journals.

**Figure 2:**
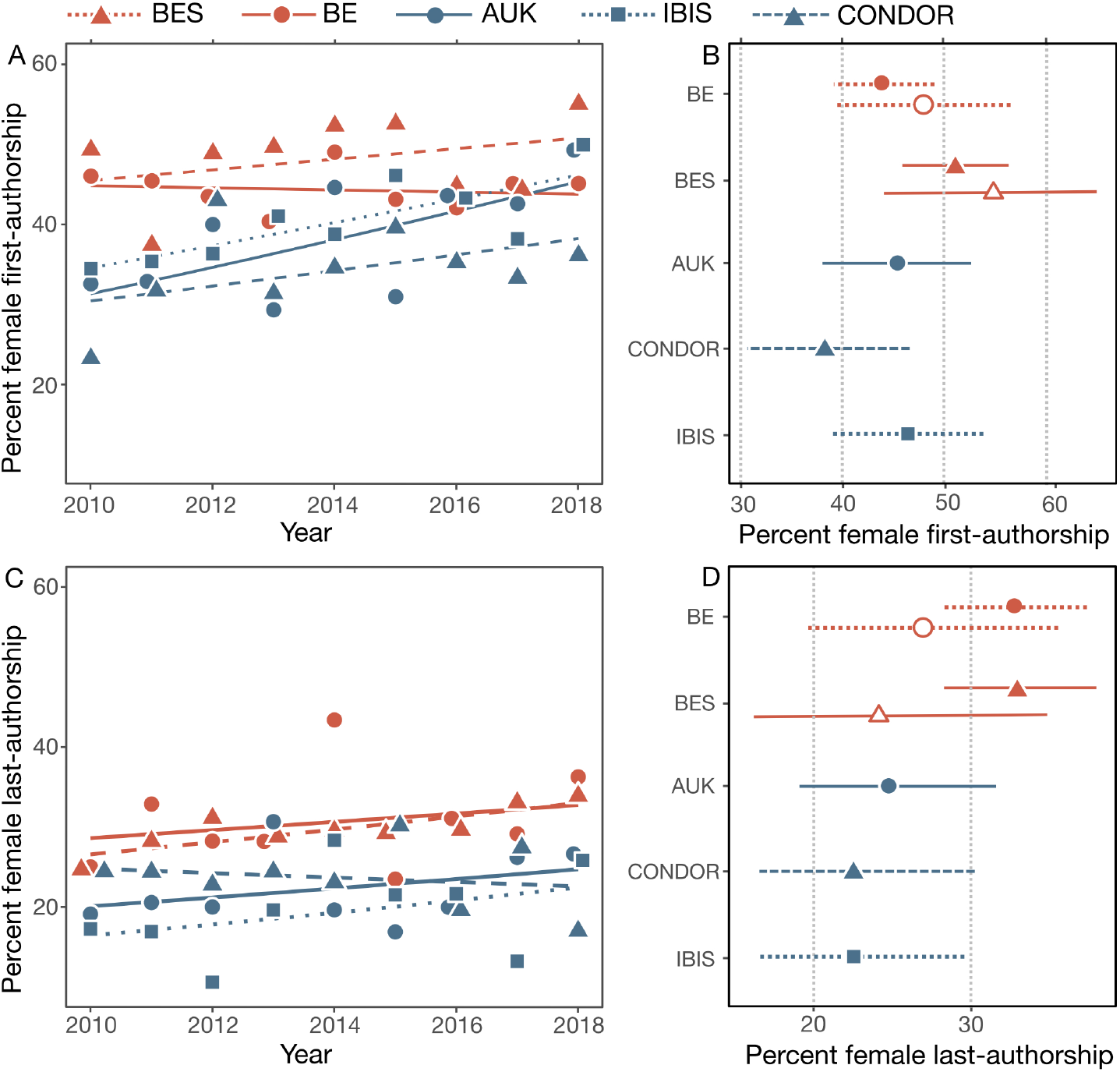
Female first- and last-authorships in 5 journals from 2010 to 2018. Female first- and last-authorships calculated as the percent of total first- or last-authorships in multi-authored papers (A, C). Percentage of female first- and last-authorships in 2018 (B, D) for each journal as well as for bird papers in BE and BES (open symbols). Percentages were calculated as marginal means of the models shown in Table 2. Papers with ambiguous authorships are not included. See Table 1 for sample sizes.

### Single-authorship papers

The percentage of single-authored papers that had female authorship changed across years in all journals, but the rate of change varied significantly (year*journal, Table 2). While the double-blind-reviewing BE initially had the most female single-authorships, that percentage declined significantly from 2010 to 2018 (–15% per year [–26, –3]), while every single-blind journal increased (BES 21% [2, 46], CONDOR 18% [–19, 78], IBIS 17% [–22, 77]) or remained constant (AUK 0% [–24, 31]; Fig. 3).

In contrast, for papers about birds, there was no significant variation in the percentage of single-authored papers having a female authorship across journals or years (Table 2, Fig. S6).

**Figure 3:**
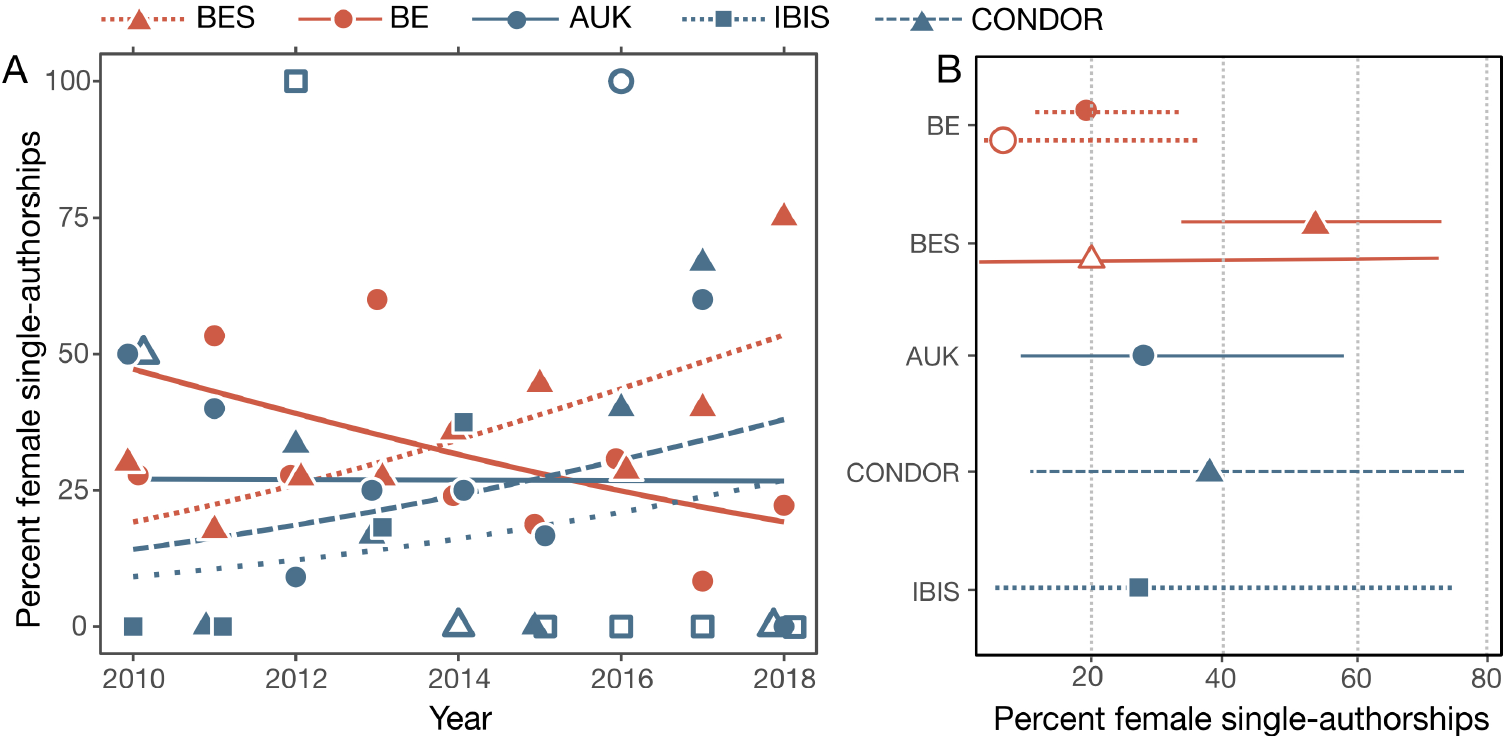
Female single-authorships in 5 journals from 2010 to 2018. (A) Percentages of single-authored papers that had female authorships in each journal, with binomial trendlines. Open shapes for years with <3 single-authorship papers. (B) Percentage of female single-authorship papers in 2018 for each journal as well as for bird papers in BE and BES (open symbols). Percentages were calculated as marginal means of the models shown in Table 2. Papers with ambiguous authorships are not included. See Table 1 for sample sizes.

### Authorships of collaboration leaders

For papers on all topics, if the last author (assumed to be the collaboration lead) was female rather than male, the proportion of other female authorships on that paper increased significantly (Table 2) by 44% [33, 56]. For bird papers alone, the proportion of other female authorships (i.e., collaborators) significantly increased 41% [26, 58] if the last-authorship was female (Table 2).

## Discussion

Our analyses show that, from 2010-2018, the journal *Behavioral Ecology* (BE) which mandates a double-blind peer review did not have higher rates female authorship than the subject-comparable *Behavioral Ecology and Sociobiology* (BES), with single-blind review. Instead, we found a general increase in the frequency of female authorship across all journals, except for single author papers, where female authorship actually decreased in the double-blind journal while increasing in the single-blind journals. Although we found fewer female (mean 35%) than male (mean 65%) authorships, only 22.8% % of 1990-2011 papers on JSTOR about ecology and evolution were written by women, suggesting that these rates reflect the gender ratios of the field, rather than a publishing bias. Overall, we find no evidence that double-blind peer review increases the incidence of female authorship. With female authorships increasing over the most recent decade in these journals, there appears to be no longer a gender bias in publication that can be attributed to the reviewing process.

The three ornithology journals had lower rates of female authorship than the more taxon-general behavioral ecology journals (BE and BES). This discrepancy does not seem to be a bias in the field of ornithology resulting from different gender ratios among ornithological authors or from a reviewer bias among ornithologists as the pattern holds even with bird papers published in the behavioral ecology journals (Fig. S3-S6). Moreover, the lower rates of female authorship in the bird journals are not likely to be due to differences in the nationalities of authorship, as IBIS, BE, and BES all publish many papers by authors outside North America. Instead, we suggest that lower rates of female authorship in the ornithology journals may be a result of women being less likely to submit to these taxon-specific ornithology journals, for some as yet unknown reason. As women in other fields tend to have broader research programs and specialize less often (Leahey, 2006), journals with more general readership might be more appealing to female scientists than the specialized ornithology journals. Alternatively, BE and BES were both founded relatively recently (1990 and 1976 respectively) with both men and women on the editorial boards and have always aimed for gender parity. In sharp contrast, the ornithology journals were all founded by small groups of men in the mid-to-late 1800s, and never had female editors-in-chief (0/19 for AUK, 0/14 for CONDOR, though the newly-appointed EIC is a woman). This awareness of potential bias may well have benefited the gender ratios of authors in BE and BES. For example, conference organizers achieve gender parity when they explicitly consider gender when inviting speakers, while those who do not do this tend to under-invite women, relative to the proportion of female society members (Débarre, Rode & Ugelvig, 2018).

In our dataset, 11% of papers had at least one authorship whose gender we could not be certain of from their first name alone. Manuscript reviewers, however, often have prior knowledge of an author’s gender, particularly amongst those with established careers and particularly in small fields such as ornithology. Reviewers may also look up unfamiliar authors to get a sense of who they are reviewing. We did not categorize the genders of ambiguous authors by using other criteria but as they represent only 11% of our sample of papers, they are unlikely to influence our findings.

While double-blind reviewing does not appear to have influenced any gender bias in publication success over the past decade in the journals that we surveyed, gender bias in academia is still a serious concern. Indeed, only ~25% of last authorships were female, in stark contrast to the 60% female undergraduate population (Eddy, Brownell & Wenderoth, 2014). This disparity between gender ratios in incoming undergraduates and tenured professors is commonly attributed to lingering effects of historical inequality. However, the disparity is larger than that predicted by that factor alone, with women less likely than men to pursue academic careers following graduate school (Shaw & Stanton, 2012). In part, this may be due to unconscious biases within academia. When asked to evaluate an application for lab manager, science faculty rated male applicants as more competent and hireable than a female applicant with an identical record, and men were offered higher starting salaries and more mentorship opportunities (Moss-racusin et al., 2012). Female scientists also tend to have fewer large, international collaborations—which are more likely to result in high impact papers–than men (Abramo, D’Angelo & Murgia, 2013; Campbell et al., 2013; Uhly, Visser & Zippel, 2017). Such a bias faced by women in their day-to-day academic life may discourage women from remaining in academia.

### The Case For

Although we find no evidence that a double-blind reviewing process currently improves gender equity in publishing in the journals that we surveyed, that does not mean that double-blind reviewing is not worthwhile. Rather, double-blind reviewing may reduce the incidence of nepotism and both institutional and geographic biases. If either these factors are thought to influence the acceptance of manuscripts in the journals that we studied, they should be studied in those journals specifically, in a recent sample of journal volumes.

There is evidence, for example, that authors familiar to reviewers, either through a personal connection or prominence in the field, are more likely to have their papers or grants accepted than unfamiliar authors (Sandstrom & Hallsten, 2008; Okike et al., 2016). In Sweden, success rates for medical grants were ~15% higher when the grant committee members were personally affiliated with the applicant (Sandstrom & Hallsten, 2008). Similarly, work by authors from prestigious universities and institutions was more likely to be successful than that of their unknown counterparts (Ross et al., 2006; Okike et al., 2016). Presumably, well-known authors from prestigious universities arrived at this level of prominence by being exceptionally good researchers and submit high-quality work. If this was the case, these authors would have high acceptance rates, whether their name and affiliations were attached to their submissions or not. However, when personal identifiers were removed, their success rates dropped 10-15% (Ross et al., 2006; Okike et al., 2016), again suggesting a strong bias in favor of the well-known.

There is also evidence from other general surveys that there are often strong geographic biases with authors from the USA, Canada, and the UK being substantially more likely to have their work accepted for publication than authors from other countries (Link, 1998; Tregenza, 2002; Ross et al., 2006; Primack & Marrs, 2008; Primack et al., 2009). Furthermore, only 2-4% of Indian and Chinese papers submitted to Biological Conservation were accepted from 2004-2007 (Primack & Marrs, 2008). As this apparent bias may be due the disadvantage of being a non-native English speaker submitting to an English journal (Tregenza, 2002; Ross et al., 2006), double-blind review may not increase acceptance rates substantially. Nonetheless, acceptance rates vary dramatically between non-English countries (Primack & Marrs, 2008), suggesting possible geographic biases which may be corrected via double-blind review.

One common criticism of double-blind review, particularly in small fields of study, is that reviewers can identify authors from the study system or location. One study, however, found that even though reviewers, especially experts in the field, attempt to guess the authors of manuscripts that they are reviewing, they are wrong 74-90% of the time (Goues et al., 2017). In ornithology, in particular, and behavioral ecology, in general, we would expect reviewers to have a higher success rate as study organisms, study sites and methods of analysis are often strongly associated with particular authors throughout their careers.

### The Case Against

For the five journals that we surveyed, the most obvious reason to avoid double-blind reviewing is that that procedure does not influence the publication rate of women scientists—and may even be detrimental (Fig. 1-3). Our analyses of two very comparable journals (BE and BES) suggest that publications by women are currently less likely to appear in the double-blind-reviewing BE. There is no obvious reason for this difference and it may simply reflect a preference for women to submit manuscripts to the journal that does not have double-blind reviews (BES). Thus any costs involved in double-blind reviewing do not seem to produce any positive benefits to female scientists submitting their papers to BE.

Several previous studies have outlined three obvious arguments against double blind reviews. First, the process of preparing a manuscript for double blind review is time-consuming if done well. Time spent removing authors’ names, and any telling details of study location, study species, references, acknowledgments, and funding, might be more profitably be spent checking statistical details, improving graph quality, or preparing data and statistical code for an online repository, all of which might be more beneficial than double blind reviews. Second, the double-blind reviewing process requires some additional editorial time if done well, checking submitted manuscripts thoroughly and corresponding with authors who have not met the journal’s requirements. This is an additional burden that might discourage authors or increase the costs of journal editing.

Finally, double-blind reviewing deprives potential reviewers of useful information when deciding whether to accept a request to review. Scientists might also be reluctant to provide additional reviews to papers that they have rejected with prejudice from a different journal, or by authors whose work they do not trust and would not be willing to review if the authors were revealed. Analogous to one of the core principles of Bayesian statistics, informative prior knowledge might well benefit the reviewing process.

We also wonder whether authors might derive some intangible and long-term benefits when reviewers know who they are. As scientists become more experienced and prominent in their field, they are likely to do more reviews, and those reviews often constitute an increasing proportion of the papers that experienced scientists read thoroughly. For many reviewers, knowledge about the quality, creativity, and relevance of research (and the researchers) is acquired in large measure from the reviewing process. Double blind reviewing thus deprives authors of that potentially important source of information. We have not seen this issue mentioned in previous studies of gender bias and double blind reviewing, and suggest it might be worth further investigation, as difficult as it might be to quantify.

### Recommendations

In our experience, journal editorial boards have strong opinions about the value of double-blind reviewing, but we hope that our analyses might help to inform those opinions. Because we are behavioral ecologists, we would advocate a cost-benefit approach to decision making. If the goal is simply to maximize what we have characterized as the benefits to double blind review, then, of course, double blind is likely to be the best course of action, unless it actually discourages author submissions. But if the goal is to maximize the net benefits, then the decision is not so clear, and some thoughtful analysis of the costs might prove informative.

## Acknowledgements

We thank the Natural Sciences and Engineering Research Council of Canada and Queen’s University for funding; Fran Bonier for useful discussion; and xxx for comments on the manuscript.

## References

Abramo G, D’Angelo CA, Murgia G. 2013. Gender differences in research collaboration. Journal of Informetrics 7:811–822. DOI: 10.1016/j.joi.2013.07.002.

Borsuk RM, Aarssen LW, Budden AE, Koricheva J, Leimu R, Tregenza T, Lortie CJ. 2009. To name or not to name: the effect of changing author gender on peer review. BioScience 59:985–989. DOI: 10.1525/bio.2009.59.11.10.

Budden AE, Tregenza T, Aarssen LW, Koricheva J, Leimu R, Lortie CJ. 2008. Double-blind review favours increased representation of female authors. Trends in Ecology and Evolution 23:4–6. DOI: 10.1016/j.tree.2007.07.008.

Campbell LG, Mehtani S, Dozier ME, Rinehart J. 2013. Gender-heterogeneous working groups produce higher quality science. PLoS ONE 8:1–6. DOI: 10.1371/journal.pone.0079147.

Cho AH, Johnson SA, Schuman CE, Adler JM, Gonzalez O, Graves SJ, Huebner JR, Marchant DB, Rifai SW, Skinner I, Bruna EM. 2014. Women are underrepresented on the editorial boards of journals in environmental biology and natural resource management. PeerJ 2:e542. DOI: 10.7717/peerj.542.

Débarre F, Rode NO, Ugelvig L V. 2018. Gender equity at scientific events. Evolution Letters 2:148–158. DOI: 10.1002/evl3.49.

Eddy SL, Brownell SE, Wenderoth MP. 2014. Gender gaps in achievement and participation in multiple introductory biology classrooms. CBE—Life Sciences Education 13:478–492. DOI: 10.1187/cbe.13-10-0204.

Engqvist L, Frommen JG. 2008. Double-blind peer review and gender publication bias. Animal Behaviour 76:e1–e2. DOI: 10.1016/j.anbehav.2008.05.023.

Fox CW, Ritchey JP, Paine CET. 2018. Patterns of authorship in ecology and evolution: First, last, and corresponding authorship vary with gender and geography. Ecology and Evolution:in press. DOI: 10.1002/ece3.4584.

Gilbert JR, Williams ES, Lundberg GD. 1994. Is there gender bias in JAMA’s peer review process? Journal of the American Medical Association 272:139–142. DOI: 10.1001/jama.1994.03520020065018.

Goues C Le, Brun Y, Apel S, Berger E, Kurshid S, Smaragdakis Y. 2017. Effectiveness of anonymization in double-blind review. arXiv preprint:arXiv:1709.01609.

Knobloch-Westerwick S, Glynn CJ, Huge M. 2013. The Matilda effect in science communication: an experiment on gender bias in publication quality perceptions and collaboration interest. Science Communication 35:603–625. DOI: 10.1177/1075547012472684.

Larivière V, Ni C, Gingras Y, Cronin B, Sugimoto CR. 2013. Bibliometrics: global gender disparities in science. Nature 504:211–213. DOI: 10.1038/504211a.

Leahey E. 2006. Gender differences in productivity: Research specialization as a missing link. Gender and Society 20:754–780. DOI: 10.1177/0891243206293030.

Link AM. 1998. US and non-US submissions. JAMA 280:246. DOI: 10.1001/jama.280.3.246.

Moss-racusin CA, Dovidio JF, Brescoll VL, Graham MJ, Handelsman J. 2012. Science faculty’s subtle gender biases favor male students. Proceedings of the National Academy of Sciences 109:16474–16479. DOI: 10.1073/pnas.1211286109.

Okike K, Hug KT, Kocker MS, Leopold SS. 2016. Single-blind vs double-blind peer review in the setting of author prestige. Journal of the American Medical Association 316:1315–1316. DOI: 10.1001/jama.2016.11014.

Primack RB, Ellwood E, Miller-rushing AJ, Marrs R, Mulligan A. 2009. Do gender, nationality, or academic age affect review decisions? An analysis of submissions to the journal Biological Conservation. Biological Conservation 142:2415–2418. DOI: 10.1016/j.biocon.2009.06.021.

Primack RB, Marrs R. 2008. Bias in the review process. Biological Conservation 141:2919–2920. DOI: 10.1016/j.biocon.2008.09.016.

Ross JS, Gross CP, Desai MM, Hong Y, Grant AO, Daniels SR, Hachinski VC, Gibbons RJ, Gardner TJ, Krumholz HM. 2006. Effect of blinded peer review on abstract acceptance. JAMA 295:1675. DOI: 10.1001/jama.295.14.1675.

Sandstrom U, Hallsten M. 2008. Persistent nepotism in peer-review. Scientometrics 74:175–189. DOI: 10.1007/s11192-008-0211-3.

Shaw AK, Stanton DE. 2012. Leaks in the pipeline: separating demographic inertia from ongoing gender differences in academia. Proceedings of the Royal Society B 279:3736–3741. DOI: 10.1098/rspb.2012.0822.

Tomkins A, Zhang M, Heavlin WD. 2017. Reviewer bias in single-versus double-blind peer review. Proceedings of the National Academy of Sciences 114:12708–12713. DOI: 10.1073/pnas.1707323114.

Tregenza T. 2002. Gender bias in the refereeing process? Trends in Ecology and Evolution 17:349–350. DOI: 10.1016/S0169-5347(02)02545-4.

Uhly KM, Visser LM, Zippel KS. 2017. Gendered patterns in international research collaborations in academia. Studies in Higher Education 42:760–782. DOI: 10.1080/03075079.2015.1072151.

Webb TJ, Hara BO, Freckleton RP. 2008. Does double-blind review benefit female authors? Trends in Ecology and Evolution 23:2006–2008. DOI: 10.1016/j.tree.2008.04.004.

